# Stimulus visibility controls the balance between attention-induced changes in contrast appearance and decision bias

**DOI:** 10.1101/091421

**Authors:** Sirawaj Itthipuripat, Kai-Yu Chang, Ashley Bong, John T Serences

## Abstract

While attention is known to improve information processing, whether attention can alter visual appearance has been a cornerstone of debate for 100+ years. Although recent studies suggest that attention can alter appearance, it has been argued that the reported appearance changes reflect response bias. Here, we provide a resolution to this debate by showing that attention has different effects on appearance and response bias depending on stimulus visibility. In a contrast judgment task where the contrast of attended and unattended stimuli varied across a full range of contrast values, human participants exhibited a substantial amount of response bias to the attended stimulus, when stimuli were hard to see. However, when stimuli were easier to see, response bias decreased and attention primarily increased perceived contrast. These results help constrain philosophical arguments about the cognitive penetrability of perception and reconcile the long-standing debate about the attention effect on appearance and response bias.

## Introduction

Despite the wealth of studies showing that attention facilitates behavioral performance by enhancing the efficiency of sensory information processing (Anton-Erxleben & Carrasco, 2013; Carrasco, 2011; Itthipuripat & Serences, 2015; Serences & Kastner, 2014; Sprague, Saproo, & Serences, 2015), a contentious debate about whether attention can actually alter the subjective visual experience has continued for over 100 years ( Anton-Erxleben, Abrams, & Carrasco, 2010, 2011; Beck & Schneider, 2016; Block, 2007, 2010; Fodor, 1984; Helmholtz, 1866; James, 1890; Ling & Carrasco, 2007; Prinzmetal et al., 1996; Pylyshyn, 1999; Raftopoulos, 2001; Schneider & Komlos, 2008; Schneider, 2011).

Recently, Carrasco, Ling, and Read introduced a psychophysical attention-cueing paradigm that measures the perceived contrast of attended and unattended visual stimuli (Carrasco, Ling, & Read, 2004). They and others have demonstrated that an attended stimulus appears to have a higher contrast than an unattended stimulus (Anton-Erxleben et al., 2010, 2011; Carrasco et al., 2004; Liu, Abrams, & Carrasco, 2009; Störmer, McDonald, & Hillyard, 2009). Using a similar method, other studies have demonstrated that the effect of attention on appearance generalizes to other stimulus features including spatial frequency (Gobell & Carrasco, 2005), object shape and size (Anton-Erxleben, Henrich, & Treue, 2007; Fortenbaugh, Prinzmetal, & Robertson, 2011), and facial attractiveness (Störmer & Alvarez, 2016). However, in another set of studies, Schneider and collegues argue that these changes in appearance are instead related to response bias induced by the attention cues (Beck & Schneider, 2016; Schneider & Komlos, 2008; Schneider, 2006, 2011). In support of this view, other studies have shown that attention can induce a response bias in a way that makes observers more likely to respond to a cued attended stimulus compared to an un-cued unattended stimulus (Luo & Maunsell, 2015; Müller & Findlay, 1987; Wyart, Nobre, & Summerfield, 2012). Moreover, unlike attention-induced changes in appearance and perceptual sensitivity (Hillyard, Vogel, & Luck, 1998; Itthipuripat & Serences, 2015; Itthipuripat, Ester, Deering, & Serences, 2014; Itthipuripat, Garcia, Rungratsameetaweemana, Sprague, & Serences, 2014; Mangun & Hillyard, 1988; Martínez-Trujillo & Treue, 2002; Reynolds, Pasternak, & Desimone, 2000; Störmer et al., 2009), attention-induced response bias does not involve selective neural modulations in early sensory cortex (Luo & Maunsell, 2015).

Together, these findings suggest support for both opposing views regarding the impact of attention on appearance. Here we reconcile these competing accounts by proposing that attention-induced changes in subjective appearance and response bias coexist and they are differentially expressed at different levels of stimulus visibility (i.e. contrast). We based our hypothesis on the assumption that when stimulus visibility is low, sensory-evoked responses are not strong enough to interact with attention signals. Thus, subjects’ judgments about stimulus appearance will be heavily influenced by response bias. On the other hand, as stimulus visibility increases, larger stimulus-evoked responses should strongly interact with attention signals in early visual cortex and give rise to changes in visual appearance.

To test this account, we modified the comparative contrast judgment paradigm developed by Carrasco and colleagues (Carrasco et al., 2004) by adding a stimulus contrast manipulation where the contrast values of the attended and unattended stimuli varied independently (Figures 1a-b; see Methods). Over all, we found that when subjects viewed low contrast stimuli, attention induced a significant response bias but minimal changes in perceived contrast. However, when subjects viewed higher contrast stimuli, response bias decreased and attention primarily increased perceived contrast.

**Figure 1.**
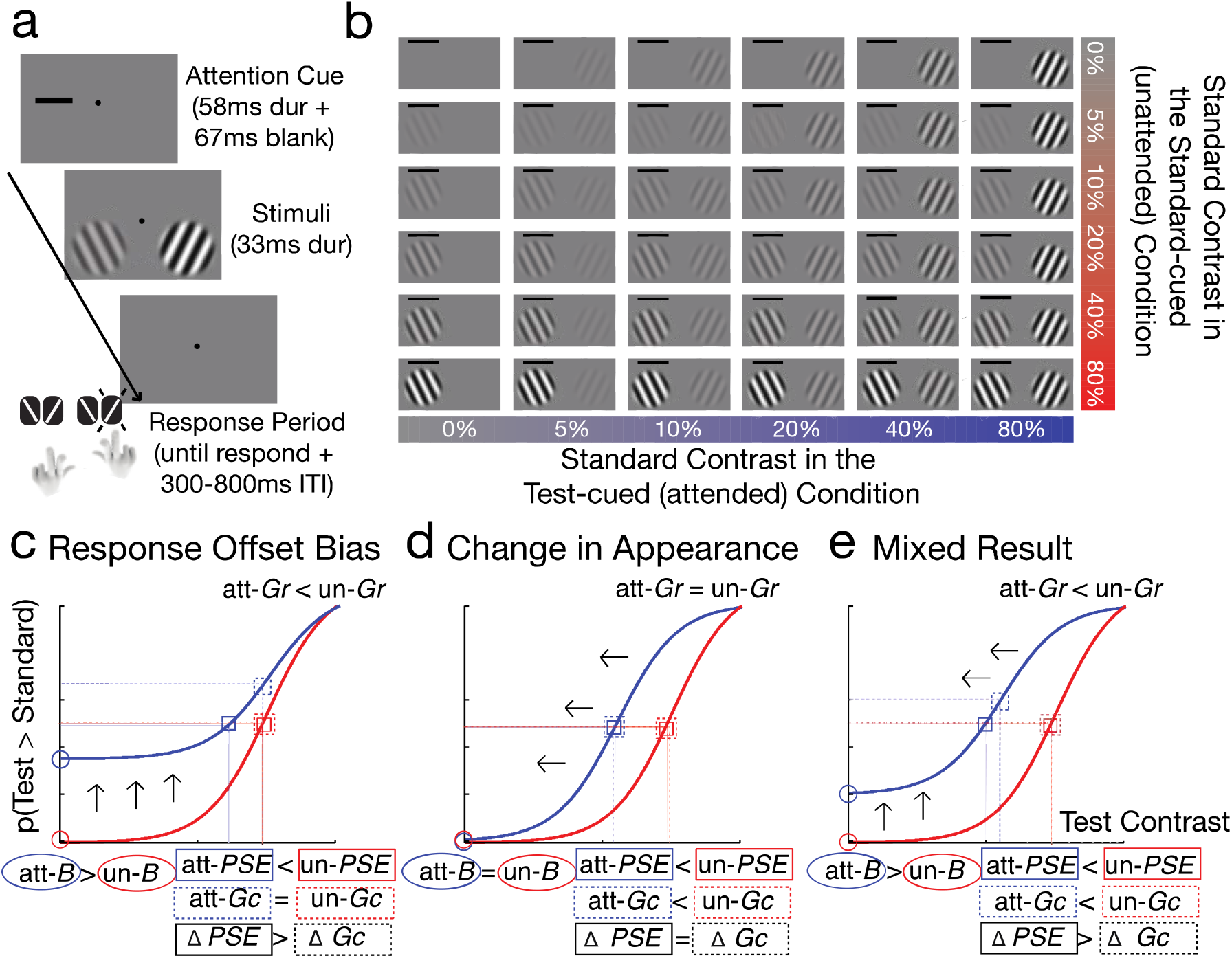
(a) A comparative contrast judgment task. Each trial started with an exogenous attention cue, followed by a set of two Gabor stimuli of which contrast values were manipulated independently from 0%-80%. Subjects reported the orientation of the stimulus (CW or CCW) that was perceived with higher contrast with the hand corresponding to the spatial location of the selected stimulus. (b) All experimental conditions showing that the contrast values of cued and uncued stimuli were manipulated independently. (c-d) Possible patterns of attentional modulations of psychometric functions. (c) Attention induces response baseline offset bias (*B;* additive upward shift) without changing contrast sensitivity (*G_c_*; horizontal shift). In this scenario, even though there is no change in *G_c_*, changes in the point of subjective quality (*PSE*; the point at which the probability value reach 50%) will be observed, which is simply due to changes in *B.* This indicates that *PSE* is not an accurate measure of contrast sensitivity when taking possible changes in *B* into account. (d) Attention alters perceived contrast of visual stimuli. In this scenario, attention should only decrease *G_c_*, but it should not change *B*, which indexes response bias driven by the attention cues. When there is no change in *B*, the *PSE* will change in a similar degree as *G_c_*. (e) Attention could induce both changes in subjective appearance indexed by *G_c_* and response bias as indexed by *B*. In this scenario, PSE will also overestimate changes in contrast sensitivity due to changes in *B*. Since the probability value cannot exceed 1, the response gain parameter (*G_r_*; the slope of the psychometric function) will decrease if *B* increases (d-e).

## Methods

### Participants

Ten neurologically healthy human observers with normal or corrected-to-normal vision were recruited from the University of California, San Diego (UCSD). All participants provided written informed consent as required by the local Institutional Review Board at UCSD. Subjects were compensated at a rate of $10 per hour. The data from one subject was excluded from the analysis because the orientation discrimination performance was below chance, leaving the data from nine subjects in the main analysis (7 female, 20-24 years old, 1 left-handed). The sample size is within the typical range for studies using similar multisession approaches(Anton-Erxleben et al., 2010, 2007; Cutrone, Heeger, & Carrasco, 2014; Sirawaj Itthipuripat, Garcia, et al., 2014; Ling & Carrasco, 2007).

### Stimuli and task

Stimuli were presented on a PC running Windows XP using MATLAB (Mathworks Inc., Natick, MA) and the Psychophysics Toolbox (Brainard, 1997; Pelli, 1997). Participants were seated 60 cm from the CRT monitor (which had a grey background of 34.51 cd/m2, 120Hz refresh rate) in a sound-attenuated and dark room.

Participants performed a comparative contrast judgment task (Figures 1 a-b), where they reported which of simultaneously presented left and right Gabor stimuli (spatial frequency = 0.33°/cycle, standard deviation of the Gaussian envelop = 2.18°, eccentricity = 13.74°) were rendered in a higher contrast value, while maintaining central fixation during the entire experiment. To examine the impact of covert spatial attention on subjects’ report about stimulus contrast, we presented an exogenous cue (0.36 ° x3.63° length x thickness) of 58ms duration (either left or right above the stimulus location) 133ms before the onset of the two Gabor stimuli (1:1 left-cued: right-cued). The contrast values of the two stimuli were independently manipulated and drawn from six contrast levels (0%, 5%, 10%, 20%, 40%, and 80% Michelson contrasts). Task difficulty was also manipulated by varying the offset values of the Gabor orientation. Here, we directly assayed response bias by including trials with 0% contrast stimuli (i.e., stimulus-absent trials), as we reasoned that any behavioral responses in favor of a cued 0%-contrast stimulus (Figures 1c and 1e) must be driven by cue-induced response bias and could not be driven by an interaction between cue-driven attention signals and the response evoked by the (absent) stimulus. Trial orders were pseudo-randomized. The participants were instructed to report if the stimulus on the left or the right visual field had a higher contrast and were specifically asked to ignore the presence of the cue. If the stimulus on the left had a higher contrast, they reported the orientation of the left stimulus (clockwise [CW] or counter-clockwise [CCW]) by pressing one of the two buttons using their left hand. If the stimulus on the right had a higher contrast, they reported the orientation of the right stimulus (CW or CCW) by pressing one of the other two buttons using their right hand. There was no response deadline and the duration of the inter-trial interval was pseudo-randomized drawn from the uniform distribution of 300-800ms. Each participant performed this task for four days. Each day contained 1,296 trials and last about 1.5 hours (5,184 trials and ~6 hours in total for each participant).

### Analysis

For each of the stimulus of interest (termed here as the standard stimulus), we calculated the probability that subjects reported the paired stimulus (termed here as the test stimulus) as higher in contrast, and we only included trials where subjects report the correct orientation of the stimulus. Since there was no stimulus presented in the 0%-contrast stimulus condition, the direction of orientation offset (CW or CCW) of individual 0%-contrast stimuli were randomly labeled before subjects performed the experiment. Correct responses to the 0%-contrast stimulus conditions determined by whether given responses matched the previously assigned labels. The difference between probability values where the cued and un-cued test stimuli were absent were used to determine whether by chance an exogenous cue has an impact on subjects’ bias to choose the cued stimulus over the uncued stimulus, even though the stimulus that was chosen was absent.

Next, we fit individual subject data with the Naka-Rushton equation using MATLAB fminserch:

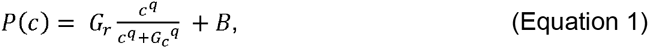

where *P(c)* is the probability the participants reported the test stimulus having a higher contrast than the standard stimulus of a given contrast value, *B* is the baseline offset (indexing response bias), *G_c_* is a contrast gain factor that controls the horizontal shift of the curve (indexing perceived appearance), *G_r_* is a multiplicative response gain factor that controls the vertical shift of the psychometric curve, and q is the exponent fixed at 2 (Carandini & Heeger, 2012). This fitting method was done separately for each standard contrast, attention condition, and orientation offset. Then, the data were collapsed across all different orientation offsets because this manipulation had little effect on the outcome (to see the data for each orientation offset condition, look in Figures S1–S2). Note that changes in subjective appearance have been traditionally indexed by changes in the point of subjective equality (*PSE;* the point at which the probability value reaches 50%) (K. Anton-Erxleben et al., 2011; Katharina Anton-Erxleben et al., 2010; Carrasco et al., 2004; Liu et al., 2009; Störmer et al., 2009). However, most past studies have not considered the possibility of changes in the baseline offset of the probability function that could be driven by response bias. Accordingly, reported *PSE* changes could be driven in part by changes in *B* (Figures 1c and 1e). Thus, we focused our primary analysis of subjective appearance on the *G_c_* parameter because it provides a more stable contrast sensitivity measure, as it is estimated simultaneously with *B* and thus takes possible changes in *B* into account. However, we also reported the traditional *PSE* results for comparison. We used three-way repeated-measures ANOVAs to test the main effects of attention conditions (test-cued/standard-cued), contrast values of standard stimuli, and orientation offsets (easy/medium/hard), and interactions between these factors on fitting parameters including *B*, *G_r_*, *G_c_,* and *PSE*. Note that we did not include the fitting data, specifically the *G_r_*, *G_c_*, and *PSE* parameters for the standard stimulus of 80% contrast because there were no test stimuli that were rendered higher than 80% contrast hence our inability to obtain the entire psychometric functions that yielded good estimations of these fitting parameters. For all fitting parameters (*B*, *G_r_*, *G_c_*, and *PSE*), post-hoc t-tests were used to examine differences between attention conditions (test-cued/standard-cued) for each standard contrast. The false discovery rate (FDR) method was used to correct for multiple comparisons.

## Results

Figure 2 shows the probability of subjects reporting the test stimulus as higher in contrast than a standard stimulus rendered at individual contrast levels (left to right columns). The blue and red circles and curves correspond to the test-cued (attended) and standard-cued (unattended) conditions, respectively. The baseline fit parameters (*B*) corresponding are shown in Figure 3a. Collapsed across attention conditions, we observed that *B* increased as the standard contrast decreased (main effect of contrast: F(5, 40) = 67.87, p < 0.001, η^2^_p_ =0.90). This finding indicates that when the standard stimulus becomes less visible, subjects are more likely to report the 0%- contrast test stimulus as having a higher contrast than the standard stimulus. We next compared the data across attention conditions. We observed that attended stimuli were associated with a higher *B* compared to unattended stimuli (main effect of attention: F(1, 8) = 26.33; p < 0.001; η^2^_p_ =0.77). Since there was actually no test stimulus present at the baseline offset of the psychometric function, this pattern of results is consistent with cue-induced response bias (Figures 1c-1d). However, the magnitude of attention effects on *B* decreased as the contrast of the standard increased (interaction between attention and contrast: F(5, 40) = 19.72, p < 0.001; η^2^_p_ =0.71). Post-hoc t-tests revealed that attention increased *B* only when the standard stimuli had low to medium contrasts (0%, 5%, 10%, and 20% with t(8)'s = 6.19, 4.04, 4.08, and 3.40; 95% confident intervals [CIs] = [0.17 0.36], [0.09 0.34], [0.06 0.23], and [0.01 0.04]; p's <0.001, = 0.004, =0.004 and =0.009, respectively; FDR-corrected threshold of 0.009) but not when they had higher contrast values (40% and 80% with t(8)'s = 1.28 and -0.82; 95% CIs = [-0.01 0.03] and [- 0.02 0.01]; p's = 0.24 and 0.43, respectively). Collectively, these results suggest that attention induces response bias when the contrast of the standard is low to medium (0%-20% contrast) but not when the contrast of the standard is higher (40%-80% contrast).

**Figure 2.**
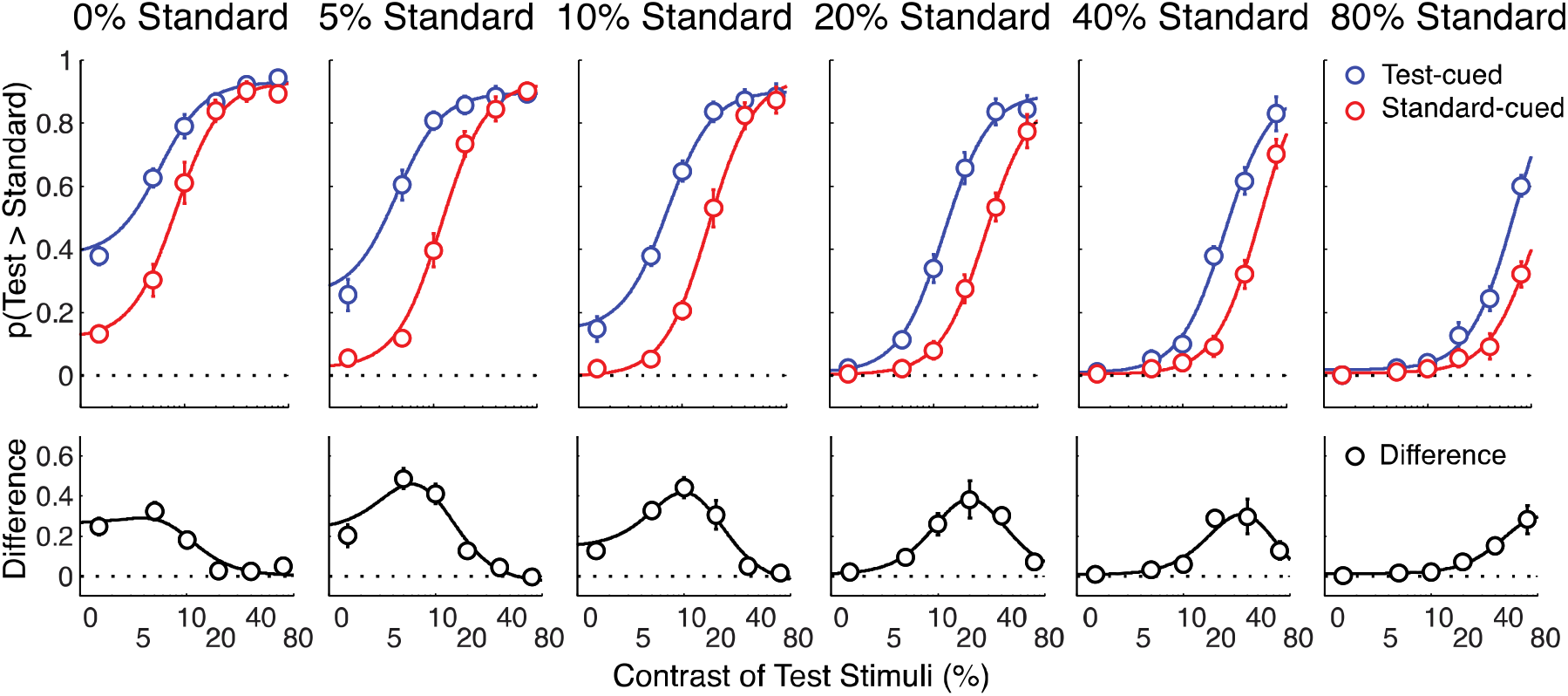
The probability of subjects reporting the test stimulus as higher in contrast than the standard stimulus, plotted as a function of the contrast of the standard (0%-80%). This probability is plotted separately for trials in which the test stimulus was cued (blue) and for trials in which the standard stimulus was cued (red). Bottom panels show the differences between the test-cued and standard-cued conditions (black). Error bars represent between-subject ±1 standard error of the mean (S.E.M).

**Figure 3.**
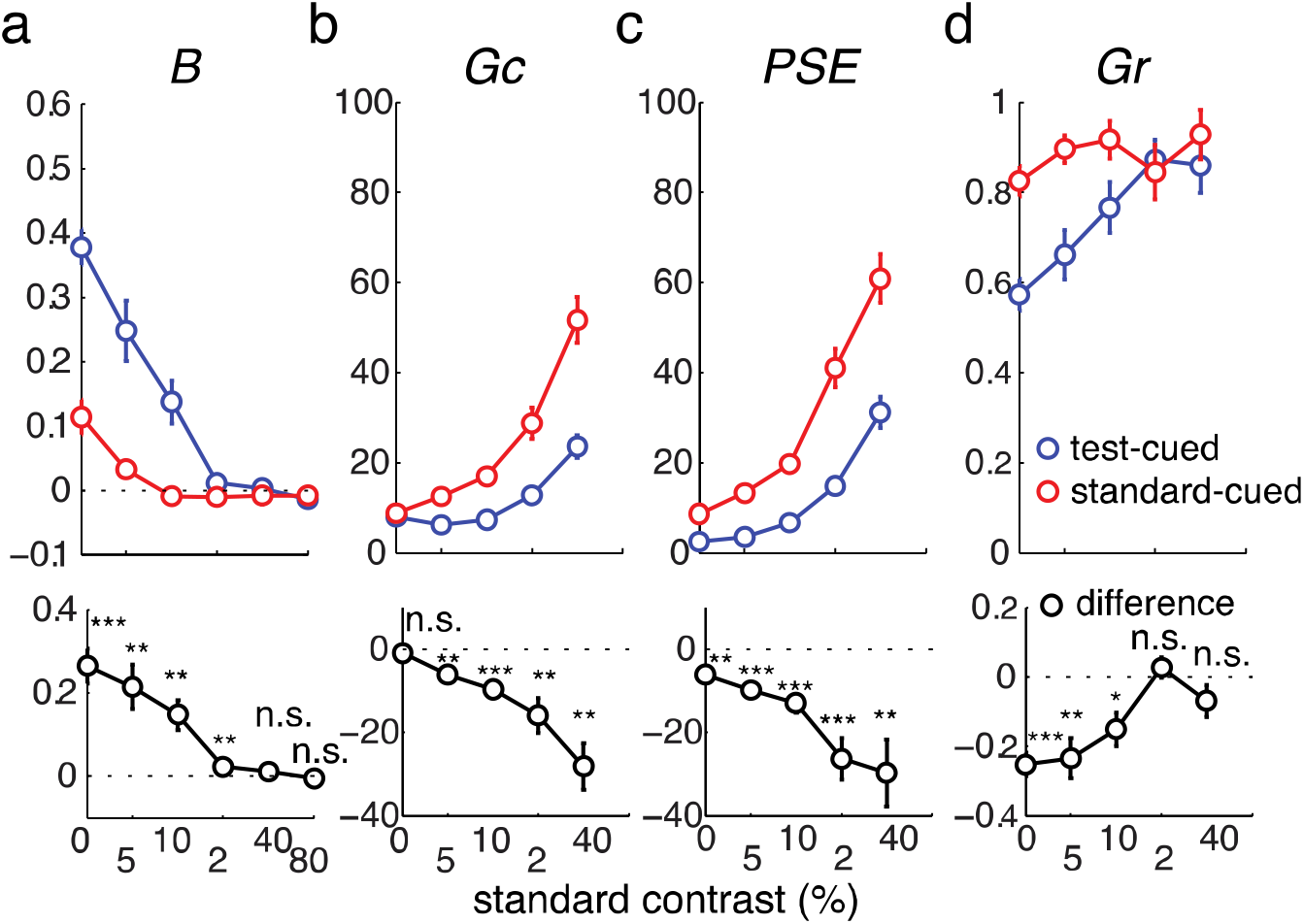
Corresponding best fit parameters for psychometric functions shown in Figure 2. *, **, and *** indicate differences in fitting parameters compared to zero with p's <0.05, < 0.01, and <0.001, respectively. Error bars represent between-subject ±1 S.E.M. Note that we did not include the *G_c_*, *PSE*, and *G_r_* parameters when the standard stimulus had a contrast of 80% because we could not obtain the entire psychometric function in this condition (see Figure 2).

For the contrast gain parameter (*G_c_*), which indexes changes in perceived contrast, we found that it increased as the contrast of the standard increased (main effect of standard contrast: F(4, 32) = 105.08; p <0.001; η^2^_p_ = 0.93). This *G_c_* shift demonstrates that when the standard stimuli became more visible, subjects were less likely to report the test stimuli as higher contrast. In contrast, attention cues had the opposite effect and led to a *G_c_* decrease and a corresponding shift of the psychometric curves to the left (main effect of attention: F(1, 8)=35.58; p <0.001; 2= 0.82). This leftward shift indicates that the attended stimuli were more likely to be reported as higher contrast compared to the unattended stimuli. Moreover, the magnitude of these attention-induced *G_c_* changes increased as a function of the contrast of the standard (interaction between attention and standard contrast: F(4, 32)=12.91, p < 0.001; 2=0.62). Post-hoc t-tests demonstrated that attention did not induce any *G_c_* change when the standard stimulus was rendered at 0% contrast (t(8) = -0.93; 95% CI = [-2.75 1.17]; p = 0.380) but that it did have an effect at all the other contrast levels (5%, 10%, 20%, and 40% with t(8)'s = -4.77, -5.35, -3.82, and -5.00; 95% CIs = [- 9.22 -3.20], [-13.84 -5.50], [-25.60 -6.33], and [-41.07 -15.13]; p's = 0.001, <0.001, = 0.005, and =0.001, respectively; FDR-corrected threshold of 0.005). Together, these results demonstrate an increasing impact of attention on perceived contrast as stimulus visibility increases. In addition, the observation that attention cues did not impact *G_c_* when the standard stimulus was rendered at 0% contrast rules out the unlikely possibility that the presentation of a cue induced a false perception of an actual stimulus. If this had occurred, *G_c_* should have shifted even when the standard was absent.

While our analysis of attention-induced response bias and perceived contrast focuses primarily on *B* and *G_c_*, respectively, we also report modulations of *PSE* and *G_r_* for completeness. The overall pattern of *PSE* modulations closely tracked the pattern of *G_c_* modulations. We found that *PSE* values increased as the contrast of the standard increased (main effect of standard contrast: F(4, 32) = 315.41; p <0.001; η^2^_p_=0.98) and decreased with attention (main effect of attention: F(1, 8)=28.51; p <0.001; η^2^_p_= 0.78; Figure 3c). The magnitude of attention-induced *PSE* changes also increased as a function of standard contrast (interaction between attention and standard contrast: F(4, 32)=9.21, p < 0.001). However, in contrast to the *G_c_* results, attention had a significant impact on the *PSE* even when the standard stimulus was absent (0% standard; t(8) = -3.91; 95% CI = [-9.72 -2.50]; p = 0.005; 2=0.54). Moreover, the attention-induced changes in *PSE* were larger than *G_c_* changes for all conditions where there were attention-induced changes in the *B* parameter (0%, 5%, 10%, and 20%; with t(8)'s = -2.83, -1.99, -3.78 and -2.43; 95% CIs = [-9.66 - 1.00], [-7.81 0.58], [-5.30 -1.28], and [-20.16 -0.53]; p's = 0.022, 0.082, 0.005, 0.041; FDR-corrected threshold of 0.005. However, for the 40%-contrast condition, where there was no attention-induced *B* change, the attentional modulations of *PSE* and *G_c_* were comparable (t(8) =- 0.20; 95% CI = [-20.06 16.85]; p =0.846). These results suggest that when there was a significant impact of attention on response bias (0%-20% standard contrast), *PSE* overestimated changes in perceived contrast compared to the contrast gain parameter (*G_c_*; Figures 1c and 1e).

For *G_r_*, note that changes in the baseline parameter (*B*) will necessarily impact *G_r_* because the value of p(test>standard) cannot exceed one. Thus, as *B* increases, *G_r_* will decrease, providing a complementary means of assessing response bias. Collapsed across attentional cue conditions, we observed a significant main effect of standard contrast (0%-40% contrast: F(4, 32) = 10.02, p < 0.001; η^2^_p_ =0.56) and attention on *G_r_* (F(1, 8) = 22.52; p < 0.001; η^2^_p_ =0.74), as well as a significant interaction between these two factors (F(4, 32) = 9.39, p < 0.001; η^2^_p_ =0.54; Figure 3d). Post-hoc t-tests revealed that this interaction was driven by an attention-related decrease in *G_r_* at the lowest thee contrast levels (0%, 5%, and 10% with t(8)'s = -7.46, -4.02, and -3.13; 95% CIs = [-0.33 -0.17], [-0.37 -0.10], and [-0.26 -0.04]; p's <0.001, =0.004, and =0.014, respectively; FDR-corrected threshold of 0.014). In contrast, the effects of attention on *G_r_* were negligible at higher contrast values (20% and 40% contrast: t(8)'s = 0.92 and -1.48; 95% CIs = [-0.04 0.10] and [-0.18 0.04x]; p's = 0.383 and 0.177, respectively). This overall pattern is consistent with the observation that attention increased *B* when the standard stimulus had a 0%-20% contrast and had little impact on *B* when the standard was higher contrast.

## Discussion

The debate about whether our subjective perceptual experience is influenced by higher cognitive functions such as attention is a contentious issue in neuroscience, psychology and philosophy (Anton-Erxleben et al., 2010, 2011; Beck & Schneider, 2016; Block, 2010; Carrasco et al., 2004; Firestone & Scholl, 2015; Fodor, 1984; Helmholtz, 1866; James, 1890; Ling & Carrasco, 2007; Prinzmetal et al., 1996; Pylyshyn, 1999; Raftopoulos, 2001; Schneider & Komlos, 2008; Schneider, 2011). Here, we show that the effects of spatial attention on visual appearance and on response bias can coexist and that the balance between these two types of modulation depends upon stimulus visibility. Specifically, we found that when the standard was low contrast, exogenous attention cues induced substantial response bias that led to an increase in the probability that subjects would choose the cued stimulus compared to the uncued stimulus, even when there was no stimulus present at the cued location (as reflected by changes in the baseline offset or *B*). However, as the contrast of the standard increased, the magnitude of response bias decreased. In contrast, when the contrast of the standard was high, spatial attention primarily changed stimulus appearance (as reflected by changes in the contrast sensitivity measure, or *G_c_*). Overall, the present findings help reconcile the long-standing debate about the effects of attention on perceptual experience and response bias, and suggest that stimulus visibility regulates the balance between these two types of attentional modulation.

Indeed, some previous studies have examined attention effects on perceived contrast of the test stimuli relative to the standard stimuli across different contrast levels: 16-36% contrast (Anton-Erxleben et al., 2011), 8% and 22% contrast (Carrasco et al., 2004), 5%-80% contrast (Cutrone et al., 2014). However, these studies only used the *PSE* metric to index changes in perceived contrast and they did not examine the possibility that the baseline parameters of the probability functions increased due to cue-related response bias. Thus, using only the *PSE* to measure contrast appearance is problematic because it can yield inaccurate estimates of changes in perceived contrast, especially when the standard stimuli are low-to-middle contrast (e.g. 0%-20% contrast in the present study; also see Figures 1c and 1d). Indeed, the data reported by Carrasco Ling, and Read suggest the possible presence of baseline changes(Carrasco et al., 2004). Moreover, these baseline shifts seemed to occur irrespective of the type of comparative judgment task that was used (i.e., reporting which stimulus has higher contrast or which stimulus has lower contrast, which hypothetically should be less affected by cue-induced response bias) (Carrasco et al., 2004). In the present study, we showed that when the standard stimulus had 0% contrast (i.e., the stimulus was absent), attention induced a substantial response bias as reflected by a large increase in the baseline parameter (*B*), yet the sensitivity of contrast perception did not changes as reflected in the contrast gain parameter (*G_c_*). However, while no change in the *G_c_* parameter was observed, we found a significant shift the *PSE* value with attention even though there was no stimulus to compare. This result suggests that in conditions where there are changes in decision-bias that induce a change in the baseline parameter (i.e., low-to-medium standard contrast), changes in the *PSE* parameter may not accurately index changes in contrast appearance.

In summary, we found that stimulus visibility regulated the balance between spatial attention effects on stimulus appearance and response bias. Not only do our results help to resolve the long-standing debate regarding the impact of attention on appearance and decision-bias, but they also provide useful insights about how the intensity of the stimulus interacts with spatial attention to influence visual perception and decision-making. In turn, this has implications for how and where attention effects play out in visual cortex (Luo & Maunsell, 2015).

## Contributions

SI conceived and implemented the experiments, analyzed the data, and wrote the manuscript. KC and AB collected the data and co-wrote the manuscripts. JTS conceived and supervised the project and co-wrote the manuscript.

## Acknowledgement

This study was funded by an NIH R01-EY025872 grant to JTS, a James S. McDonnell Foundation grant to JTS, and an HHMI International Student Research Fellowship grant to SI.

## Supplementary Information

**Figure S1.**
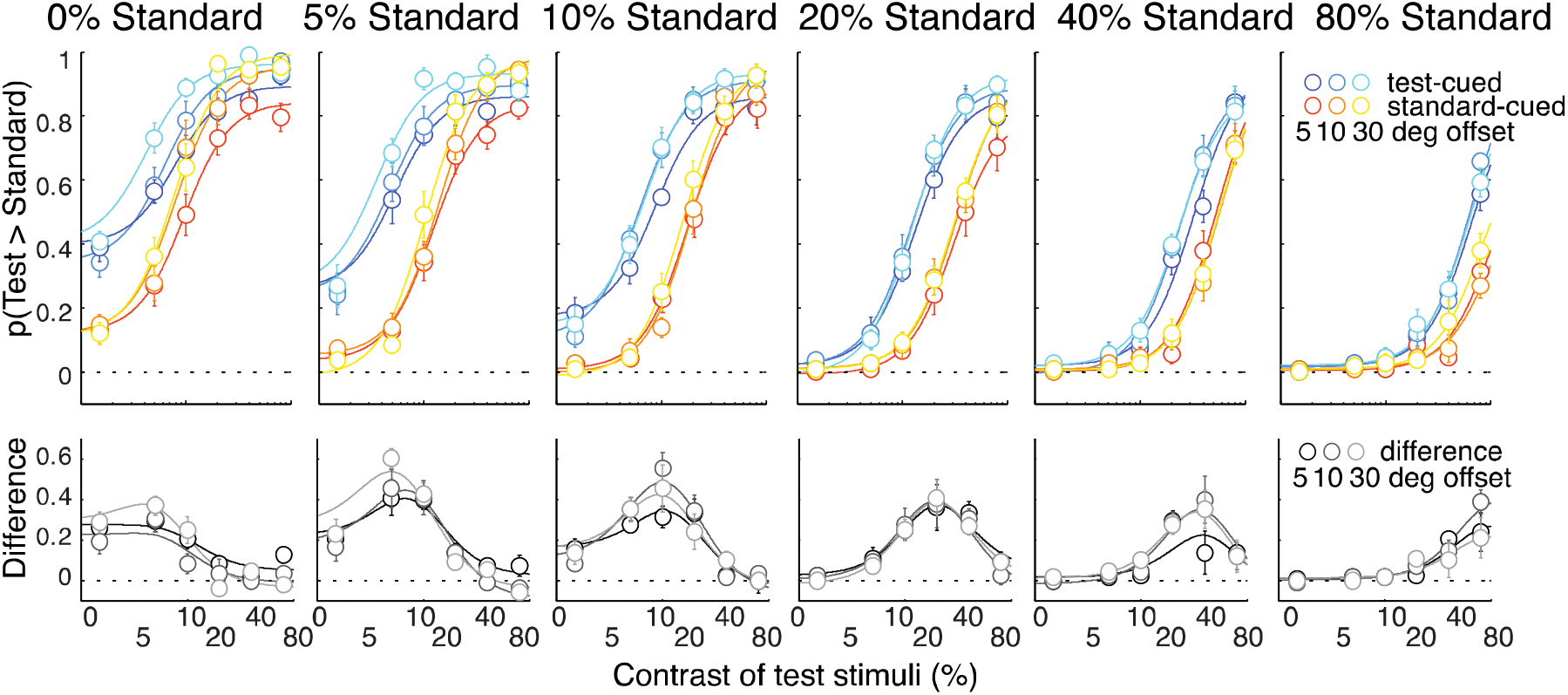
The probability of subjects reporting the test stimulus as higher in contrast than the standard stimulus, plotted as a function of the contrast of the standard (0%-80%). This probability is plotted separately for trials in which the test stimulus was cued (cold colors) and for trials in which the standard stimulus was cued (hot colors), and for trials with stimuli of different orientation offsets. Bottom panels show the differences between the test-cued and standard-cued conditions (gray-scaled colors). Error bars represent between-subject ±1 standard error of the mean (S.E.M).

**Figure S2.**
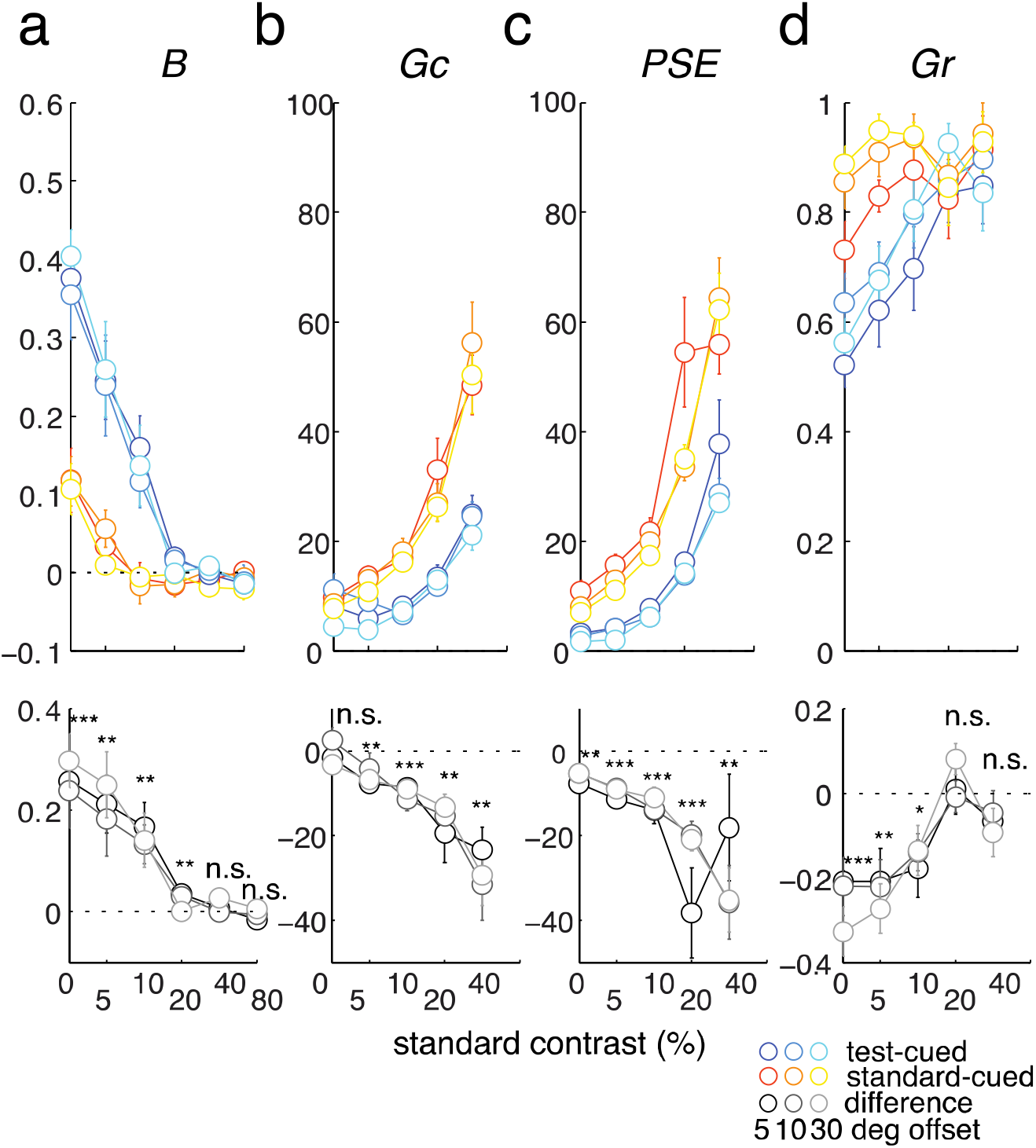
Corresponding best fit parameters for psychometric functions shown in Figure S1. *, **, and *** indicate differences in fitting parameters compared to zero with p's <0.05, < 0.01, and <0.001, respectively. Error bars represent between-subject ±1 S.E.M. Note that we did not include the *G_c_*, *PSE*, and *G_r_* parameters when the standard stimulus had a contrast of 80% because we could not obtain the entire psychometric function in this condition (see Figure S1). Note that there were a marginal main effect of difficulty on the contrast gain parameter (G_c_), and significant main effects of difficulty on the PSE and the response gain parameter (G_r_) (F(2, 16)'s = 2.58, 11.38, and 10.89, with p's = 0.107, < 0.001, = 0.001, respectively). However, task difficulty did not change the degree at which attention modulated the *G_c_*, *PSE*, and *G_r_* parameters as there was no interaction between attention and task difficulty (F(2, 16)'s ≤ 0.53; p's ≥ 0.597). In addition, there were no main effects of task difficulty or interactions between task accuracy and attention on the *B* parameter (F(2, 16)'s ≤ 0.78; p's ≥ 0.477).

